# Neuropeptide F receptor acts in the *Drosophila* prothoracic gland to regulate growth and developmental timing

**DOI:** 10.1101/2019.12.16.878967

**Authors:** Jade R. Kannangara, Michelle A. Henstridge, Linda M. Parsons, Shu Kondo, Christen K. Mirth, Coral G. Warr

## Abstract

As juvenile animals grow, their behaviour, physiology, and development need to be matched to environmental conditions to ensure they survive to adulthood. However, we know little about how behaviour and physiology are integrated with development to achieve this outcome. Neuropeptides are prime candidates for achieving this due to their well-known signalling functions in controlling many aspects of behaviour, physiology and development in response to environmental cues. In the growing *Drosophila* larva, while several neuropeptides have been shown to regulate feeding behaviour, and a handful to regulate growth, it is unclear if any of these play a global role in coordinating feeding behaviour with developmental programs. Here, we demonstrate that Neuropeptide F Receptor (NPFR), best studied as a conserved regulator of feeding behaviour from insects to mammals, also regulates development in *Drosophila*. Knocking down *NPFR* in the prothoracic gland, which produces the steroid hormone ecdysone, generates developmental delay and an extended feeding period, resulting in increased body size. We show that these effects are due to decreased ecdysone production, as these animals have reduced expression of ecdysone biosynthesis genes and lower ecdysone titres. Moreover, these phenotypes can be rescued by feeding larvae food supplemented with ecdysone. Further, we show that NPFR negatively regulates the insulin signalling pathway in the prothoracic gland to achieve these effects. Taken together, our data demonstrate that NPFR signalling plays a key role in regulating animal development and may thus play a global role in integrating feeding behaviour and development in *Drosophila*.

## INTRODUCTION

When faced with variation in the quantity and quality of the diet, young animals must adjust their feeding behaviour, metabolism, growth, and developmental time. Failure to do so has profound consequences on their ability to survive to adulthood and to resist future stress [1–3]. Extensive research into the regulation of food intake has uncovered a handful of neuropeptides that mediate changes in feeding behaviour in response to diet, including the highly conserved Neuropeptide F signalling pathway [4–7]. Developmental processes, such as growth and developmental time to adulthood, are controlled through the action of the conserved insulin and steroid hormone signalling pathways [8–10]. However, we know little about the extent to which feeding behaviour and developmental processes are coordinated, and the molecular mechanisms necessary for this coordination.

The fruit fly *Drosophila melanogaster* provides an excellent model in which to study the molecular mechanisms that integrate feeding behaviour with developmental processes. *Drosophila* development proceeds through three larval stages (instars), after which the animal initiates pupariation and metamorphosis to become an adult. The timing of the transitions between these developmental stages is regulated by a series of precisely-timed pulses of the steroid hormone, ecdysone, produced and secreted by the prothoracic gland (PG; [3]). Because these insects grow primarily during the larval stages, ecdysone dictates the length of the growth period, ceasing growth once metamorphosis begins [3, 8, 10]. In this way, ecdysone determines final adult size.

The PG produces and secretes ecdysone in response to various environmental cues, such as the day-night cycle, nutrition, and tissue damage [10–13]. These external cues are communicated to the PG via the action of a number of secreted peptides. Nutritional signals are particularly important, and are communicated throughout the body via the insulin signalling pathway. When larvae are well fed, they secrete insulin-like peptides (Dilps) into the bloodstream [14]. In the *Drosophila* PG the insulin receptor (InR) is activated by the Dilps, which in turn leads to the activation of ecdysone biosynthesis genes and therefore the production of ecdysone [8–10]. Starvation early in the third larval instar delays the onset of metamorphosis by delaying the timing of an early ecdysone pulse [8–10, 15]. Later in the third larval instar starvation accelerates developmental timing [10, 16], presumably by accelerating the production of at least one of the later ecdysone pulses. This highlights how the effects of nutrition change growth outcomes over developmental time.

As well as regulating development, the quantity and quality of nutrients in the diet also causes larvae to alter both the amount and the quality of foods they consume [17, 18]. Several peptide hormones and neuropeptides have been shown to regulate different aspects of feeding behaviour in the fly [19]. Amongst these, Neuropeptide F (NPF) signalling increases feeding rates and affects food choice in response to poor food quality [6, 19]. The mammalian homologue of NPF, Neuropeptide Y, also regulates feeding behaviour in response to food quality [20, 21]. To guarantee that the animal survives, these changes in feeding behaviour must be appropriate to the changes needed in the different stages of animal development. We therefore wondered if any of these neuropeptides act as global coordinators of development and feeding behaviour in response to nutritional signals.

Here, we show that the NPF receptor (NPFR) regulates development in *Drosophila* by regulating the production of ecdysone in the PG. We further show that NPFR signalling exerts its effects on developmental timing and body size by interacting with the insulin signalling pathway in the PG, revealing that it acts as a previously undescribed regulator of insulin signalling in this gland. Our data demonstrate that NPF signalling, well known for regulating feeding behaviour across species, also plays a key role in regulating animal development by affecting the production of developmental hormones.

## RESULTS AND DISCUSSION

### NPFR signalling regulates developmental timing

Our aim was to determine whether any of the neuropeptides that control feeding behaviour also act to alter development in response to nutrition. Because the known effects of nutrition on developmental time are controlled by ecdysone production in the PG, we hypothesised that such neuropeptides would act on receptors on the PG cells. We therefore knocked down a set of receptors for neuropeptides known to regulate feeding behaviour in larvae (Table S1) specifically in the PG. To do this we used the *phantom* (*phm*)-Gal4 driver to drive expression of RNAi constructs for the different receptors, as well as the expression of dicer II *(dcrII)* to enhance the RNAi knockdown [22].

We found that when we knocked down *NPFR* in the PG with the v9605 RNAi line, we observed a significant delay to pupariation of about 35 hours (Fig. 1A, p>0.001). With a second *NPFR* RNAi line (v107663), we only observed a significant delay when compared to the UAS-*NPFR* RNAi parental control, and not the *phm*-Gal4 parental control (Fig. 1B, p = 0.01). Therefore, to further verify that NPFR regulates developmental timing, we tested an *NPFR* loss of function mutant strain (*NPFR^SK8^*; [23]). The *NPFR^SK8^* mutant larvae also displayed a significant delay in time to pupariation compared to a heterozygous control (~15 hours; Fig. 1C, p = 0.008).

**Figure 1:**
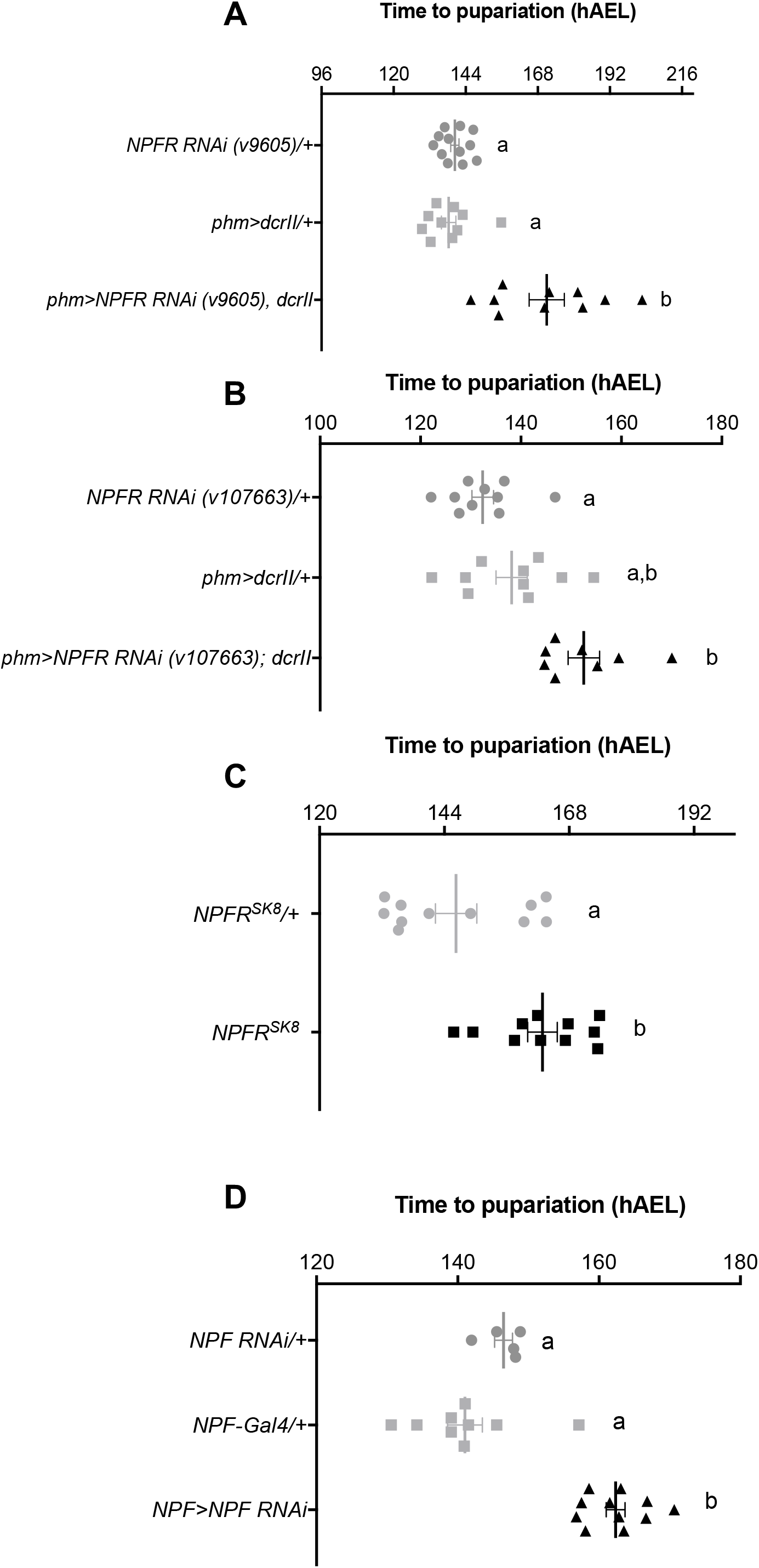
NPFR regulates developmental timing. **(A)** Knockdown of *NFPR* (using UAS-NPFR RNAi v9605) specifically in the PG (using *phm-Gal4>dcrII)* results in a significant delay in time to pupariation compared to controls (p<0.0001). **(B)** Knockdown of *NPFR* specifically in the PG using a second, independent RNAi line (UAS-NPFR RNAi v107663) also results in a significant delay in time to pupariation (p=0.0002). **(C)** *NPFR* null mutants (*NPFR^SK8^*) have a significant delay in time to pupariation (p=0.008). **(D)** Knockdown of *NPF* in NPF-expressing neurons (using *NPF*-Gal4) results in a significant developmental delay (p=0.029). hAEL= hours after egg lay. Error bars represent ±1 SEM for all graphs. In each experiment, genotypes sharing the same letter indicate that they are statistically indistinguishable from one another, while genotypes with contrasting letters indicate that they are statistically different (ANOVA and pairwise *t* tests). Each point represents a biological replicate of 15-20 animals.

NPFR could alter developmental timing because it regulates PG development. To test this, we dissected PGs from *phm-GFP>NPFR RNAi; dcrII* wandering larvae, measured their size, and examined their morphology. PG size was indistinguishable between *phm-GFP>NPFR RNAi; dcrII* and control larvae (Fig. S1, p = 0.528). Furthermore, the morphology of the PG itself appeared similar to that of the control. This data suggests that NPFR signalling does not control the development of the PG, but rather its function.

NPFR is a G-protein coupled receptor that is activated by the neuropeptide NPF [7]. To further test the role of NPFR signalling in developmental timing, we knocked down NPF specifically in the NPF-producing neurons using *NPF*-Gal4. We found that these larvae exhibited a 10-hour delay in time to pupariation when compared to controls (Fig. 1D, p = 0.029). Together, these data therefore suggest that NPF acts on NPFR on the PG cells to regulate developmental timing.

### NPFR regulates the production of ecdysone in the prothoracic gland

Given that NPFR does not seem to regulate PG development, we next tested whether NPFR signalling regulates the primary function of the PG – to produce ecdysone. We reasoned that if NPFR acts in the PG to regulate ecdysone production, then feeding *phm>NPFR RNAi; dcrII* larvae with ecdysone should rescue the developmental delay. Consistent with this prediction, supplying 20-hydroxyecdysone (20E), the active form of ecdysone, to *phm>NPFR RNAi; dcrII* larvae completely restored normal developmental timing (Fig 2A p > 0.0001). Interestingly, we found that these animals pupariate even faster than controls (Fig 2A), suggesting that they may have an increased sensitivity to ecdysone. When we quantified the ecdysone titre, we found that *phm>NPFR RNAi; dcrII* animals produced significantly less ecdysone later in the third instar, between 32 and 56 hours after the third instar moult, when compared to controls (Fig. 2B).

**Figure 2:**
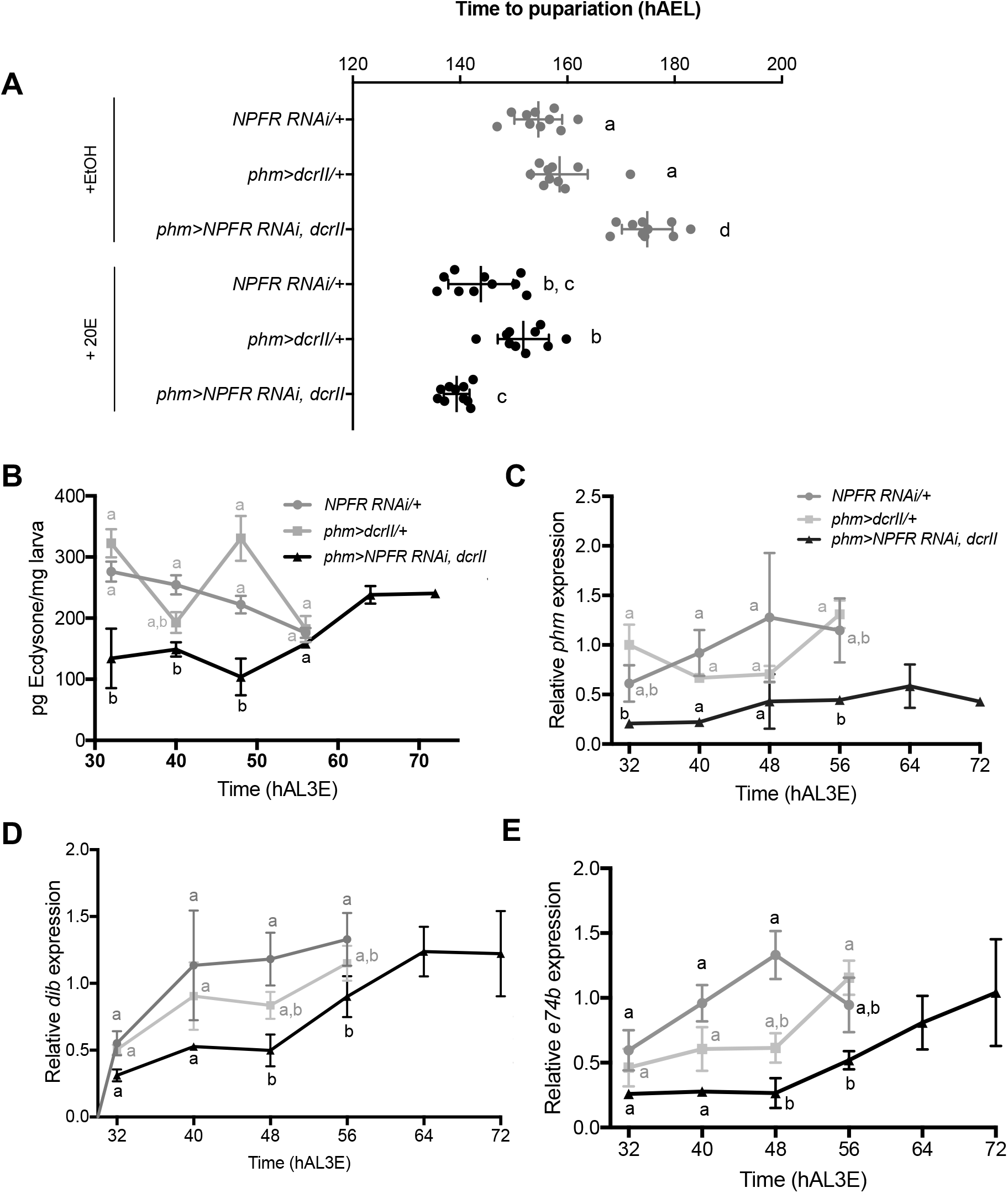
NPFR regulates the production of ecdysone in the prothoracic gland. **(A)** Time to pupariation was measured for *phm>dcrII; NPFR RNAi* (v9605) larvae fed on either food supplemented with 96% EtOH (grey) or 20-hydroxyecdysone (20E) (black). Supplying *phm>NPFR RNAi; dcrII* larvae with 20E is able to completely rescue the developmental delay seen when *NPFR* is knocked down in the PG (p<0.0001). Each point represents a biological replicate of 15-20 animals. **(B)** *phm>NPFR RNAi; dcrII* animals have an overall reduction in ecdysone titre compared to parental controls during the late third instar. hAL3E = hours after L3 ecdysis. Five biologically independent replicates of 8-10 larvae were measured for each time point. Relative expression of **(C)** *phm*, **(D)** *dib* and **(E)** *e74B* in*phm>NPFR RNAi; dcrII* animals is overall reduced as determined by quantitative PCR. Values were normalised using an internal control, *Rpl23.* hAL3E = hours after L3 ecdysis. Expression level of each gene was standardised by fixing the values at 32hrs in *NPFR* RNAi (9605)/+ as 1 in all panels. Approximately 8-15 larvae were used for each sample, and five biologically independent samples for each time point. Error bars represent ±1 SEM for all experiments. In each experiment, genotypes sharing the same letter indicate that they are statistically indistinguishable from one another, while genotypes with contrasting letters indicate that they are statistically different (ANOVA and pairwise *t* tests).

This reduction in total ecdysone concentration could be due to a defect in either its biosynthesis or in its secretion. To distinguish between these two possibilities, we quantified the expression levels of two CYP450 ecdysone biosynthetic genes, *phm* and *disembodied (dib).* The mRNA expression levels of these two enzymes are well-established as reliable proxies for ecdysone biosynthesis [9, 11, 13]. Additionally, the expression of an ecdysone response gene, *ecdysone-induced protein 74EF (e74B),* was quantified as a readout of ecdysone signalling activity. When *NPFR* was knocked down in the PG, there was an overall reduction in *phm* and *dib* between 32 and 56 hours after the third instar moult compared to controls (Fig. 2C, D). Further, *e74b* expression was reduced compared to controls (Fig. 2E), demonstrating lower levels of ecdysone signalling activity in larvae where *NPFR* was knocked down in the PG. These data suggest that NPFR signalling is involved in the regulation of ecdysone biosynthesis in the PG, although does not rule out that it could play additional roles in regulating ecdysone secretion.

### Loss of NPFR signalling phenocopies loss of insulin signalling

Changes in insulin signalling also regulate development time by regulating the rate of ecdysone synthesis [8–10, 13]. This has been demonstrated in animals which are hypomorphic for loss of insulin signalling (complete loss causes early lethality; [24]), such as flies that bear a heteroallelic combination of mutations in the *insulin receptor (InR),* or are homozygous for a loss of function mutation of the adaptor protein, *chico* [15]. These animals take longer to reach metamorphosis and have decreased adult body sizes [15, 25]. As *NPFR* mutants are homozygous viable, like *chico* mutants, we hypothesised that they may be hypomorphic for loss of insulin signalling. In support of this, we found that in addition to being developmentally delayed, *NPFR^SK8^* mutants have smaller body sizes compared to controls (Fig. 3A, p > 0.0001). Similarly, *NPF >NPF RNAi* animals are smaller than controls (Fig. 3B, p = 0.00437). To check that the body size defect was not a result of decreased food intake due to altered feeding behaviour, we quantified food intake in *NPFR^SK8^* mutants on our standard fly food. This showed that under well fed conditions there were no significant differences in the amount of food consumed compared to controls (Fig. S2A, p = 0.414).

**Figure 3:**
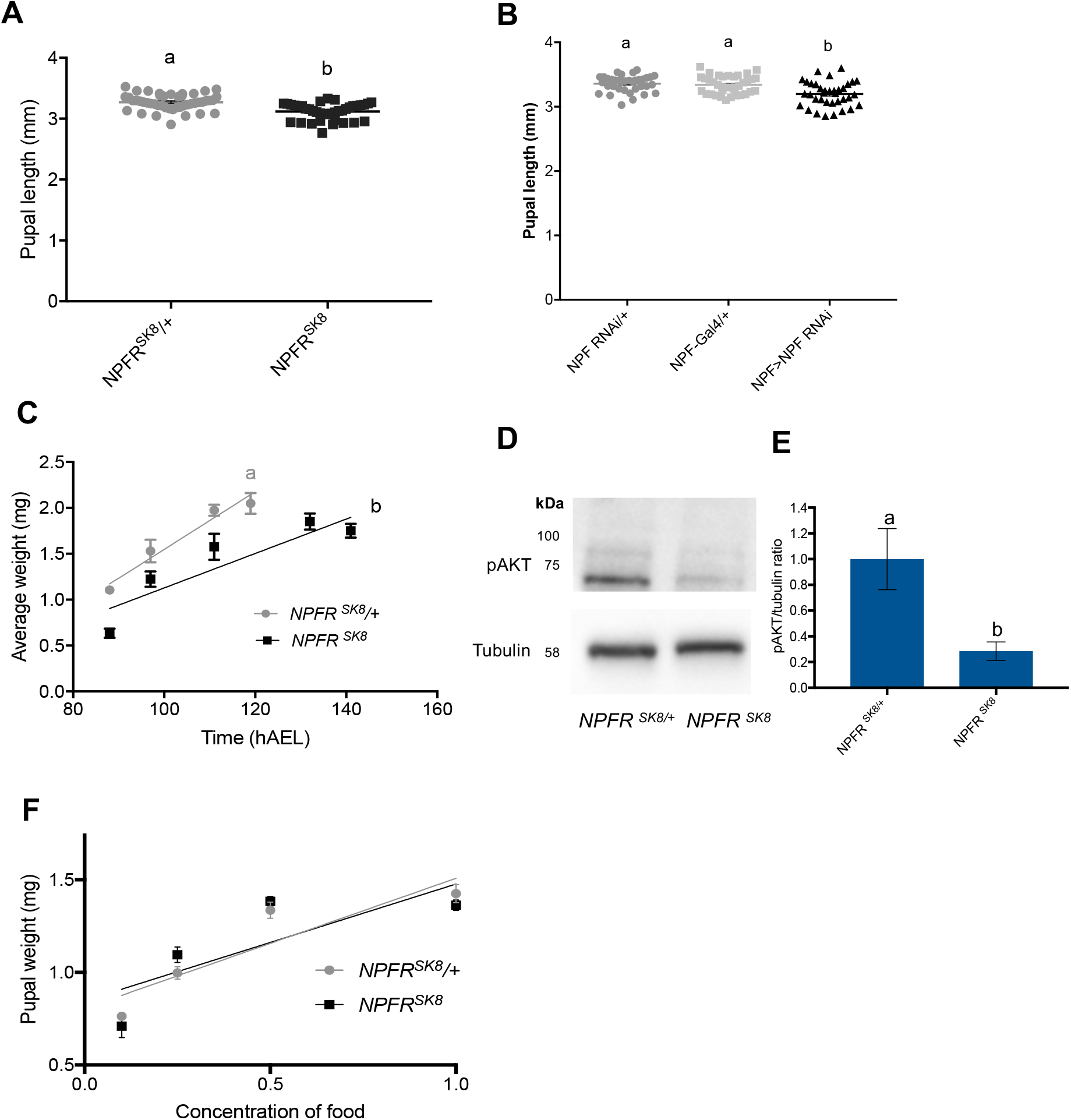
*NPFR^SK8^* mutants phenocopy loss of insulin signalling. **(A)** *NPFR^SK8^* mutants and **(B)** *NPF>NPF-RNAi* animals have a smaller body size compared to controls, as measured by pupal length (p<0.0001, p=0.00437, respectively). Each point represents an individual pupa, and no less than 40 individuals were tested per genotype. **(C)** *NPFR^SK8^* mutants have a reduced rate of growth compared to controls (p<0.01, linear regression analysis). hAEL = hours after egg lay. 10-15 animals were tested per time point for each genotype. **(D)** *NPFR^SK8^* mutants have reduced levels of phosphorylated Akt (pAkt). **(E)** Quantification of pAkt/Tubulin densities were standardised by fixing the values of *NPFR^SK8^/+* to 1. Six biological replicates of 10 animals were used per genotype. **(F)** *NPFR^SK8^* mutants fed on diets of decreasing caloric density do not adjust their body size differently to controls (p>0.05, linear regression analysis). 10 biological replicates of 10-15 animals were used per diet. For all graphs, genotypes sharing the same letter indicate that they are statistically indistinguishable from one another while genotypes with contrasting letters indicate that they are statistically different (ANOVA and pairwise *t* tests). Error bars represent ±1 SEM for all graphs.

Whole-animal mutants of the insulin signalling pathway have smaller body sizes due to reduced growth rates [26]. We therefore measured the growth rate of *NPFR^SK8^* mutants. This showed that *NPFR ^SK8^* mutants have a significantly reduced growth rate compared to controls (Fig. 3C). Together, these data suggest that animal-wide loss of NPFR phenocopies animal-wide reduction in insulin signalling.

Given these similarities in phenotype, we hypothesised that NPFR interacts with the insulin signalling pathway to regulate ecdysone production. To test this idea, we quantified insulin signalling levels in *NPFR* mutants by measuring levels of phosphorylated protein kinase B (pAKT), a downstream component of the insulin signalling pathway. A significant reduction in pAKT level was observed in *NPFR^SK8^* mutants compared to controls (Fig. 3D, E, p>0.001). This demonstrates that whole animal loss of *NPFR* leads to an overall reduction in insulin signalling.

To further explore this, we looked at another well-known phenotype of loss of insulin signalling; response to nutrition. Wild-type animals that are fed on less-nutritious foods have a reduction in final body size [27], presumably because of the resulting reduction in insulin signalling under poor nutritional conditions. When insulin signalling is suppressed in an organ, the organ loses its ability to adjust its size in response to nutrition [28]. If *NPFR* mutants have an overall reduction in insulin signalling, then they should also have decreased body size plasticity in response to less nutritious foods. We therefore fed *NPFR^SK8^* mutants diets of varied caloric concentration, and measured pupal weight as an indication of body size. This showed that these animals had the same sensitivity to nutrition as controls, with indistinguishable slopes between body size and the caloric concentration of the food between genotypes (Fig. 3F). Taken together, this data suggests that loss of *NPFR* reduces, but does not fully ablate, overall insulin signalling. The reduction in insulin signalling is sufficient to cause reduced body size and growth rate, but not enough to interfere with plasticity in body size in response to poor nutrition.

### NPFR negatively regulates insulin signalling in the prothoracic gland

Given that *NPFR^SK8^* mutants have reduced insulin signalling, and that knocking down *NPFR* in the PG reduces ecdysone synthesis, we next wanted to determine if NPFR modifies insulin signalling specifically in this organ. Changes in insulin signalling in the PG could cause a developmental delay under one of two different scenarios: 1) if insulin signalling is increased in the PG early in third instar larvae, or 2) if insulin signalling is reduced in the PG in the mid to late third instar. This is because insulin signalling has different roles before and after an important developmental checkpoint in third instar larvae, known as “critical weight” [10, 15, 29]. Prior to critical weight, either starving animals or reducing insulin signalling results in a developmental delay [10, 15, 30]. This occurs because low levels of insulin signalling delay the timing of the ecdysone pulse that is necessary to trigger the critical weight checkpoint [10]. After critical weight has been achieved, starving animals has the opposite effect and accelerates developmental timing [10, 13, 16]. This has previously been described as the “bail out response”, referring to the fact that under starvation conditions (when insulin signalling is low), the developmental program encourages the animal to pupariate [3, 31]. Thus, an increase in insulin signalling in late third instar animals should cause a developmental delay.

We therefore set out to determine if loss of NPFR in the PG reduces or increases insulin signalling in this tissue, and if this changes over the third larval instar period. To do this we examined the localisation of the transcription factor, Forkhead Box class O (FoxO), a negative regulator of insulin signalling [32], in PG cells. When insulin signalling is high FoxO is cytoplasmic, and when insulin signalling is low FoxO is localised to the nucleus [32]. We knocked down *NPFR* in the PG and determined FoxO localisation in both early and mid-late third instar animals. In both cases, PG cells in *phm> NPFR RNAi; dcrII* animals had significantly more cytoplasmic FoxO than controls (Fig 4A and 4B, p > 0.01). This suggests that insulin signalling activity is increased in the PG in both early and late third instar larvae when *NPFR* is knocked down specifically in this tissue, and thus that NPFR normally functions to repress insulin signalling in the PG in third instar larvae. The developmental delay that was observed by knocking down *NPFR* throughout the third instar could then be explained if the increase in insulin signalling pre-critical weight is not sufficient to accelerate developmental timing, but the increase in insulin signalling during the late third instar is sufficient to cause a developmental delay.

**Figure 4:**
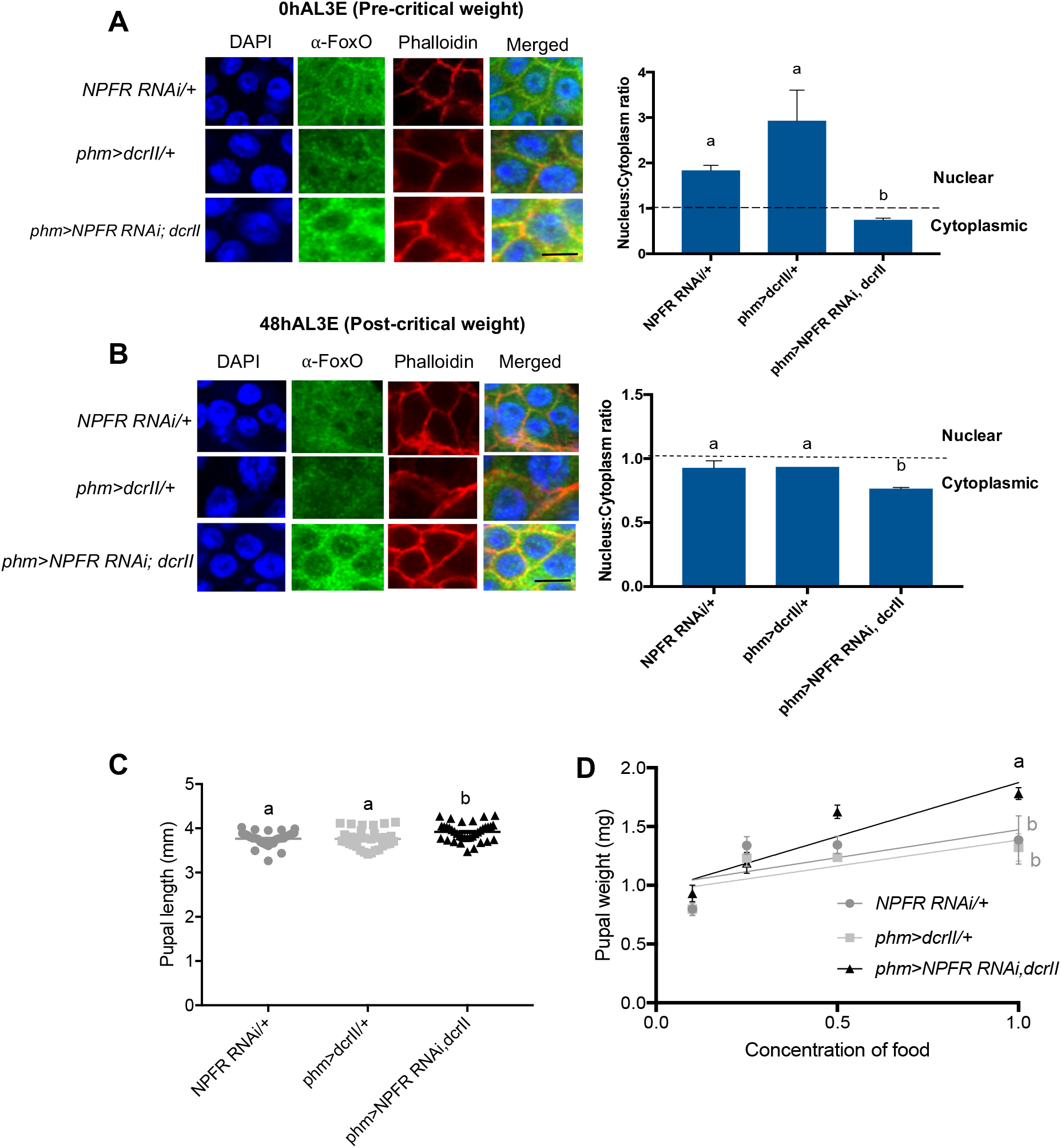
NPFR negatively regulates insulin signalling in the prothoracic gland. **(A)** Knocking down *NPFR* specifically in the PG results in increased cytoplasmic FoxO accumulation in pre-critical weight larvae (0hAL3E) compared to controls (p=0.01). **(B)** Knocking down *NPFR* specifically in the PG results in primarily cytoplasmic FoxO in post-critical weight larvae (48hAL3E) compared to controls (p=0.0011). hAL3E = hours. after L3 ecdysis. Eight – ten PGs were analysed per genotype. Scale bar is 10μm. **(C)** Knocking *NPFR* down specifically in the PG results in an increase in final body size as measured by pupal length (p=0.0239). Each point represents an individual pupa and no less than 40 individuals were tested per genotype. For **(A) – (C)**, error bars represent ±1 SEM. Genotypes sharing the same letter are statistically indistinguishable from one another, while genotypes with different letters are statistically different (ANOVA and pairwise *t* tests) **(D)** *phm>NPFR RNAi; dcrII* animals display significantly different changes in final body size compared to controls when fed on diets of decreasing caloric density (p<0.01, linear regression analysis). 10 biological replicates of 10-15 animals were assessed per diet.

We next measured body size when *NPFR* is knocked down in the PG, using pupal length as an indication of final body size. This was of interest as increasing insulin signalling before critical weight would be expected to cause a decrease in body size, whereas increasing insulin signalling after critical weight would be expected to cause an increase in body size. We found that *phm> NPFR RNAi; dcrII* animals are larger than controls (Fig. 4C, p = 0.0239). To ensure the body size alteration was not a result of increased food intake, we quantified food intake in the *phm> NPFR RNAi; dcrII* animals. This showed that the amount of food consumed was indistinguishable from controls (Fig. S2B, p = 0.588). Taken together, while the FoxO localisation data suggests NPFR is able to repress insulin signalling throughout L3, the combination of a developmental delay and increased body size observed when we knock down *NPFR* in the PG suggests that NPFR function in the PG is most important post critical weight.

Lastly, we looked at the response to nutrition. If knocking down *NPFR* in the PG causes increased insulin signalling in the gland, this could result in animals being more sensitive to nutrition [28]. To test this, we knocked down *NPFR* in the PG, and animals were fed on one of four concentrations of food. We then measured pupal weight as an indication of final body size (Fig. 4D). On a diet of standard caloric concentration (1x), *phm> NPFR RNAi; dcrII* animals were significantly larger than controls. As the calorie concentration in the food decreases, we observed a much steeper decrease in body size for *phm> NPFR RNA; dcrII* animals compared to controls. This demonstrates that body size is indeed more sensitive to nutrition in these animals, further supporting a negative regulatory role in insulin signalling.

Taken together, given that NPFR regulates insulin signalling in the PG, these data suggest that under nutritional stress NPF could be both acting on NPFR neurons in the brain to regulate feeding behaviour [6, 19] while also signalling through NPFR in the PG to regulate developmental timing. In this way, NPF signalling could act as a nexus between feeding behaviour and development.

### NPFR regulates developmental timing by acting downstream of the insulin receptor in the prothoracic gland

Finally, we conducted genetic interaction experiments to determine where in the insulin signalling pathway NPFR acts in the PG. We first overexpressed a constitutively active and ligand-independent form of InR *(InR^CA^)* in the PG. As expected [33], expression of *InR^CA^* specifically in the PG significantly reduced the time to pupariation (Fig 5A, p < 0.01). When we knocked down *NPFR* while simultaneously expressing *InR^CA^* in the PG, we observed a significant developmental delay, similar to that seen with PG-specific knockdown of *NPFR* alone (Fig. 5A, p > 0.0001). This suggests that NPFR functions downstream from InR.

**Figure 5:**
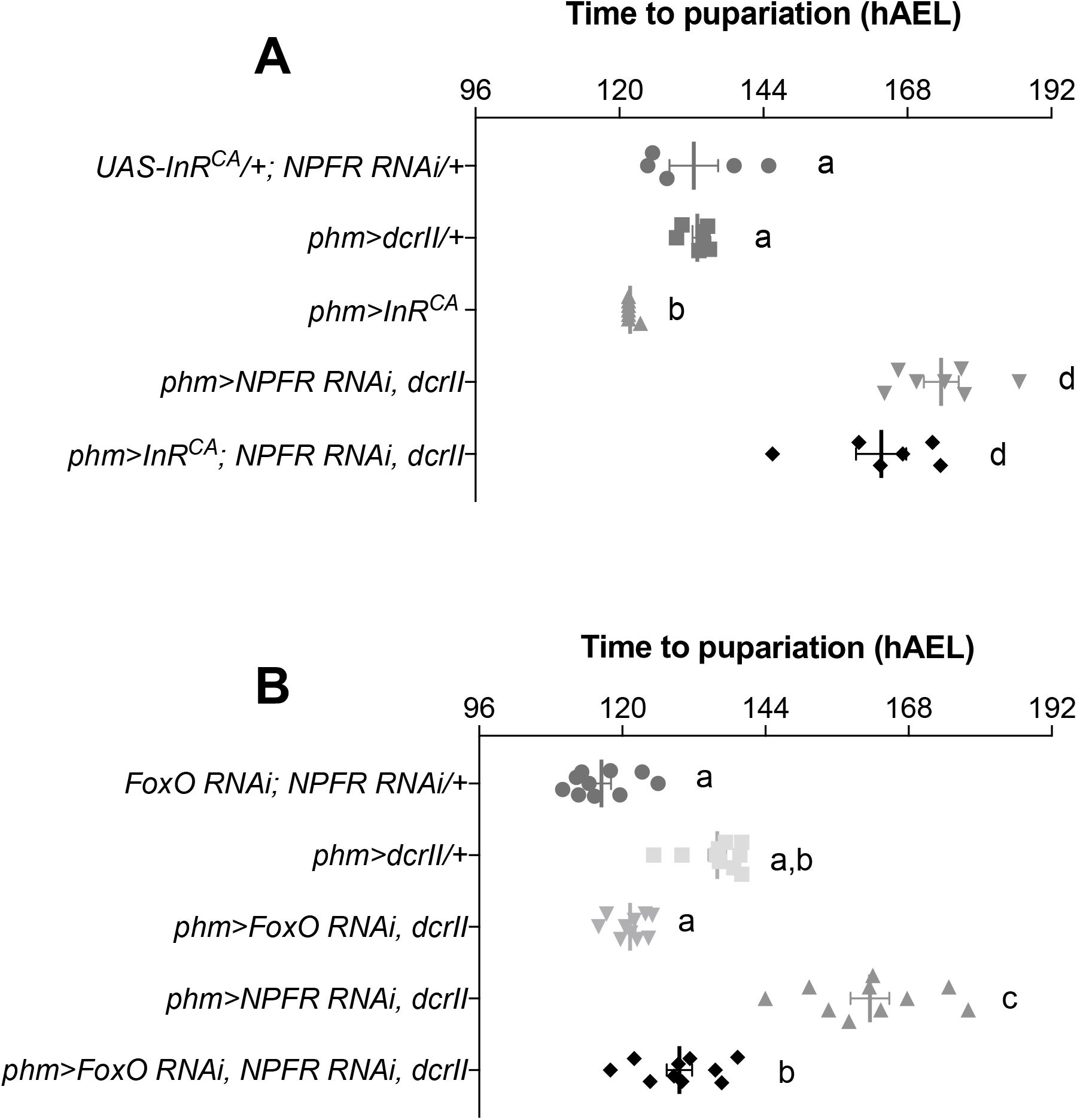
NPFR interacts with the insulin signalling pathway to control developmental timing. **(A)** Knockdown of *NFPR* while simultaneously expressing a constitutively active form of *InR*(*InR*^*CA*^) specifically in the PG results in a significant developmental delay (p<0.0001), similar to PG-specific knockdown of *NPFR* alone. **(B)** Knockdown of both *NPFR* and *FoxO* specifically in the PG partially rescues the developmental delay seen with PG-specific knockdown of *NPFR* alone (p<0.0001). hAEL= hours after egg lay. Error bars represent ±1 SEM. In each experiment, genotypes sharing the same letter indicate that they are statistically indistinguishable from one another, while genotypes with contrasting letters indicate that they are statistically different (ANOVA and pairwise *t* tests).

We then asked if NPFR acts upstream of FoxO. Mutations in *foxo* do not affect body size in fed animals [34], likely due to an unchanged development time. However, altering FoxO activity does impact developmental timing and size in starved larvae [13]. This is presumably because reducing FoxO in the PG of fed animals is insufficient to further increase insulin signalling in this gland, and so animals pupariate at normal times. If NPFR functions upstream of FoxO, then the developmental delay seen when *NPFR* is knocked down in the PG should be rescued when *foxo* is simultaneously knocked down. Therefore, we knocked down both *NPFR* and *foxo* in the PG and found that this was able to partially rescue the delay seen when knocking down alone (Fig. 5B, p > 0.0001). These results therefore suggest that NPFR functions to regulate the insulin signalling pathway downstream of InR and upstream of FoxO.

In summary, here we have described a new role for the conserved feeding regulator, NPFR, in the regulation of developmental timing, animal growth rate and body size. In the PG, our data supports a role for NPFR in negatively regulating the insulin signalling pathway and ecdysone biosynthesis. Our genetic interaction data suggests NPFR acts downstream of the insulin receptor in the PG, and perhaps plays a role in keeping the insulin signalling pathway in check to ensure that the ecdysone pulses are produced at the correct time. By contrast to the results we obtained in the PG, we found that whole animal loss of NPFR generates phenotypes resembling reduced insulin signalling. The simplest explanation of these contrasting phenotypes is that NPFR has a second role in regulating developmental timing and body size elsewhere in the fly, perhaps due to a role in regulating insulin-like peptide production or secretion. These data thus not only highlight a previously undescribed mechanism by which insulin signalling and ecdysone production are regulated in the PG, but also demonstrate how a single neuropeptide signalling pathway can have functionally diverse roles within an organism in response to the same environmental cue.

Our findings raise the strong possibility that NPF may indeed coordinate feeding behaviour and growth. How might it do so? Other peptides known to function in the regulation of ecdysone production are produced either in neurons that directly innervate the PG, such as PTTH [11], or in other tissues and secreted into circulation, such as the Dilps [14, 35]. Either a local or systemic source of NPF is therefore possible for activating NPFR in the PG. While neuropeptides such as NPF are best described as having local modes of action, in the adult fly NPF has recently been shown to be secreted from the midgut into the hemolymph, where it can act systemically [23]. In the larva NPF is known to be expressed in dopaminergic neurons in the brain, as well as cells in the midgut [20]. To our knowledge it has not been shown to be expressed in neurons that innervate the larval PG. This suggests that it is more likely that systemic rather than local NPF activates NPFR in the larval PG cells. Systemic NPF produced in response to nutritional stress could thus act on both NPFR neurons in the brain to regulate feeding behaviour and on NPFR in the PG to regulate developmental timing, and in so doing NPF could coordinate feeding behaviour and development.

In conclusion, this study has provided evidence to show that NPFR signalling, best known for its regulation of feeding behaviour, also functions in the *Drosophila* PG to control developmental timing and body size via regulation of insulin signalling and ecdysone production. To our knowledge NPF represents the first neuropeptide described to play a role in regulating both feeding behaviour and development in response to nutritional conditions, and thus first candidate for coordinating these processes in response to environmental cues. Given that the mammalian homologue of NPF, NPY, also has a role in regulating feeding behaviour in response to nutritional stress, it would be of great interest to explore if it too is a candidate for coordinating behaviour and development.

## Acknowledgements

We would like to thank Karyn Moore, Emily Kerton and the Australian *Drosophila* biomedical research facility (OzDros) for technical support; Michael O’Connor for providing fly stocks; and Pierre Leopold for the FoxO antibody. M.A.H is a National Health and Medical Research Council (NHMRC) Early Career Fellow. This work was supported by an Australian Research Council (ARC) grant to C.G.W and C.K.M.

## Author contributions

C.G.W and C.K.M conceived the experiments, interpreted the data and co-led the work. J.R.K conceived the experiments, interpreted the data, and performed the experiments. M.A.H interpreted the data and assisted with experiments. L.M.P interpreted the data. S.K. generated the NPFR mutant *Drosophila* strain. J.R.K, C.G.W and C.K.M wrote the manuscript, with assistance from L.M.P.

## MATERIALS AND METHODS

### *Drosophila* stocks

The following stocks were used: *w^1118^* (BL5905), *NPF*-Gal4 (BL25682), InR^CA^ (BL8263; a constitutively active form of InR) and UAS-FoxO RNAi (BL32993) from the Bloomington *Drosophila* Stock Centre, UAS-*NPFR* RNAi (v9605), UAS-NPFR RNAi (v107663), UAS-NPF RNAi (108772) from the Vienna *Drosophila* Resource Centre, *NPFR^SK8^* mutant (Ameku et al., 2018, Kondo and Ueda, 2013), *phm*-Gal4-22, UAS-*mCD8*.:GFP and UAS-dicerII; *phm*-Gal4-22, gifts from Michael O’Connor, University of Minnesota, Minneapolis (Ono et al., 2006). All flies were maintained at 25°C on fly media containing, per litre: 7.14 g potassium tartrate, 0.45 g calcium chloride, 4.76 g agar, 10.71g yeast, 47.62g dextrose, 23.81g raw sugar, 59.52g semolina, 7.14mL Nipagen (10% in ethanol) and 3.57mL propionic acid.

### Developmental timing assays and body size analysis

Parental flies were allowed to lay eggs on 25mm apple juice agar plates for 3-4 hours. Twenty-four hours later, 15 L1 larvae were picked into standard food vials. Ten replicates were collected from each cross. Time to pupariation of the F1 offspring were scored every 8 hours. Larvae for all experiments were raised inside an insulated, moist chamber at 25 degrees in the dark. Each set of genetic crosses included a UAS-RNAi or Gal4 control crossed to *w^1118^*; the genetic background for the RNAi library from the VDRC. As a proxy for body size, following their eclosion photos of the pupal cases from the developmental timing assays were taken using a light compound microscope at 2.5x magnification. Pupal case length was measured using Fiji.

### Immunocytochemistry

For PG morphology studies, wandering larvae from each genotype were collected, and anterior halves of the larvae were dissected and fixed for 30 minutes in 4% formaldehyde in PBTx (0.01% Triton-X in Phosphate Buffered Saline (PBS)). Samples were washed 4 times over one hour in PBTx and then incubated in 50ul RNAase for 20 minutes. Samples were incubated in DAPI (1ul in 400ul PBTx) for 2 minutes, and washed in PBTx. Samples were stored in Vectashield (Vector laboratories) and PGs were dissected under a light compound microscope in PBS. Dissected PGs were mounted onto a slide and were visualised using confocal microscopy (Olympus CV1000). Measurements of PG area were quantified using Fiji. For FoxO staining, larvae were staged at L3 and at the appropriate time points, dissected and fixed in 4% formaldehyde in PBS for 45mins at room temperature. Samples were then washed in PBTx and blocked for 30mins in 5% goat serum in PBTx and rabbit anti-FoxO (a gift from Dr. Pierre Leopold, 1:500) was added to 5% goat serum in PBTx. Samples were allowed to incubate at 4 degrees overnight. We then washed the samples in PBTx, and anti-rabbit Alexa 488 (Invitrogen, 1:500) in 5% goat serum in PBTx was added in the dark and allowed to incubate for 1.5 hours at room temperature. Samples were then washed and incubated in DAPI (1:400 PBTx) for 2 minutes. After washing in PBTx again, we added Phalloidin (1:1000) to the samples and allowed them to incubate at room temperature for 20 minutes. Samples were washed and stored in Vectashield (Vector laboratories) before further dissection onto poly-L-lysine coated coverslips and analysed using confocal microscopy.

### Growth rate

Parental flies were allowed to lay on 25mm apple juice agar plates for 3-4 hours. Parent flies were removed and the eggs were allowed to develop for a further 24 hours. 15-20 L1 larvae were picked into standard food vials and were allowed to develop for a further 72 hours. Six-eight replicates were picked for each genotype. Individual larvae were then floated in 20% sucrose to retrieve them from the vials, and one replicate was weighed using a microbalance (Mettler Toledo) each morning and evening until the larvae started to pupariate. Weight over time was recorded and analysed using Prism 7.

### Ecdysone feeding

To make 20E food, a stock solution of 10mg/ml of 20E (Cayman Chemical) was dissolved in 96% EtOH. To reach a final concentration of 0.15mg/ml, 15ul of the stock solution was added per 1g blended fly media. For the control food, 96% EtOH was used without 20E addition. Ten young L3 larvae were picked into the vials and allowed to feed *ad libitum.* Time to pupariation was measured every 8 hours. 10 replicates were used per genotype.

### Quantitative PCR (qPCR)

Total RNA was extracted from the anterior halves of 10-15 larvae using TRIsure (Bioline). After DNase treatment, total RNA concentration was quantified and no more than 5ug of total RNA was converted to cDNA using a 1:1 mix of oligo DT and random hexamer primers, and reverse transcriptase (Bioline). qPCR was performed using SYBR Green PCR MasterMix (Bioline). Primer sequences for *phm, dib* and *rpl23* were borrowed from McBrayer *et al.,* 2007. Sequences for *e74b* are as follows: *e74B* (F-5’ CGGAACATATGGAATCGCAGTG, R-5’ CATTGATTGTGGTTCCTCGCTG 3’).

### Ecdysone Titre Quantification

Larvae were synchronized by collecting newly ecdysed L3 larvae every 2 hrs. A sample of eight to ten larvae was weighed on a microbalance (Mettler Toledo) and then preserved in methanol. Prior to assaying, the samples were homogenized and centrifuged, and the resulting methanol supernatant was dried. Samples were resuspended in 50ul of enzyme immunoassay (EIA) buffer (0.4 M NaCl, 1 mM EDTA, 0.1% bovine serum albumin, and 0.01% sodium azide in 0.1M phosphate buffer). 20E EIA antiserum and 20E acetylcholinesterase tracer were purchased from Cayman Chemicals.

### Immunoblotting

Five L3 larvae were homogenised in 80μl of lysis buffer (50mM Tris-HCl (pH 7.5), 150mM NaCl, 2.5mM EDTA, 0.2% Triton X, 5% glycerol, complete EDTA-free protease inhibitor cocktail (Roche)) and spun at 500g for 5 min at 4°C. Reducing buffer was added to all samples before boiling and separation by SDS-PAGE (any kDa TGX, Biorad) followed by transfer onto an Immobilon-P membrane (Millipore). Membranes were probed with either 1:1,000 anti-phosphorylated *Drosophila* Akt (Cell Signalling, 4054S), or 1:1,000,000 anti-α-tubulin (Sigma, B-5-1-2), washed and incubated with HRP-conjugated secondary antibody (1:10,000, Southern Biotech). Immunoblots were developed using ECL prime (GE healthcare) and imaged using a chemiluminescence detector (Vilber Lourmat). pAkt blot images were quantified using Fiji and differences between genotypes determined by unpaired t-tests from six biological replicates.

### Nutritional plasticity

Food of varied caloric concentrations was made by diluting our standard food (SF) as described above with 0.5% agar (Gelita). The food concentrations used were 0.1x (10% SF, 90% agar), 0. 25x (25% SF, 75% agar), 0.5x (50% SF, 50% agar) and 1x (100% SF). Eggs were picked onto these diluted foods and pupal weight was measured using a microbalance (Mettler Toldeo). For each genotype at least ten replicates of 15 larvae were raised on each food concentration. Differences in genotypes was determined by linear regression analysis using GraphPad Prism.

### Quantification of food intake

Newly moulted third-instar larvae were transferred to freshly dyed food (4.5% blue food dye) and allowed to feed for 1 hr. After feeding, larvae were removed from food using 20% sucrose solution, washed in distilled water and dried. Replicates of 10 larvae were homogenised in 80μl of cold methanol and centrifuged for 10 min at 4°C. 60μl of supernatant from each sample was analysed in a spectrophotometer at 600nm. As standards, a two-fold dilution series of food dye, starting at a concentration of 4μl dye/ml methanol was used. Five-Six biological replicates were analysed per genotype.

