## Supplemental Table and Figures for "Neuropeptide F receptor acts in the *Drosophila* prothoracic gland to regulate growth and developmental timing"

### SUPPLEMENTARY FIGURE TITLES AND LEGENDS

**Table S1. List of neuropeptide receptors known to bind to ligands that regulate feeding behaviour in *Drosophila* larvae**

| Neuropeptide | Receptor(s) | Reference |
|---|---|---|
| Allatostatin A (AstA) | Allatostatin Receptor 1 and 2 (AstA-R1 and AstA-R2) | [36] |
| Drosulfakinin (Dsk) | Cholecystokinin-like receptor at 17D1 and Cholecystokinin-like receptor at 17D3 (CCKLR-17D1 and CCKLR-17D3) | [37] |
| Neuropeptide F | Neuropeptide F Receptor (NPFR) | [6, 19] |
| Hugin (Hug) | Pyrokinin 2 Receptor 1 and 2 (PK2-R1 and PK2-R2) | [38, 39] |
| Short neuropeptide F (sNPF) | Short neuropeptide F Receptor (sNPFR) | [40] |

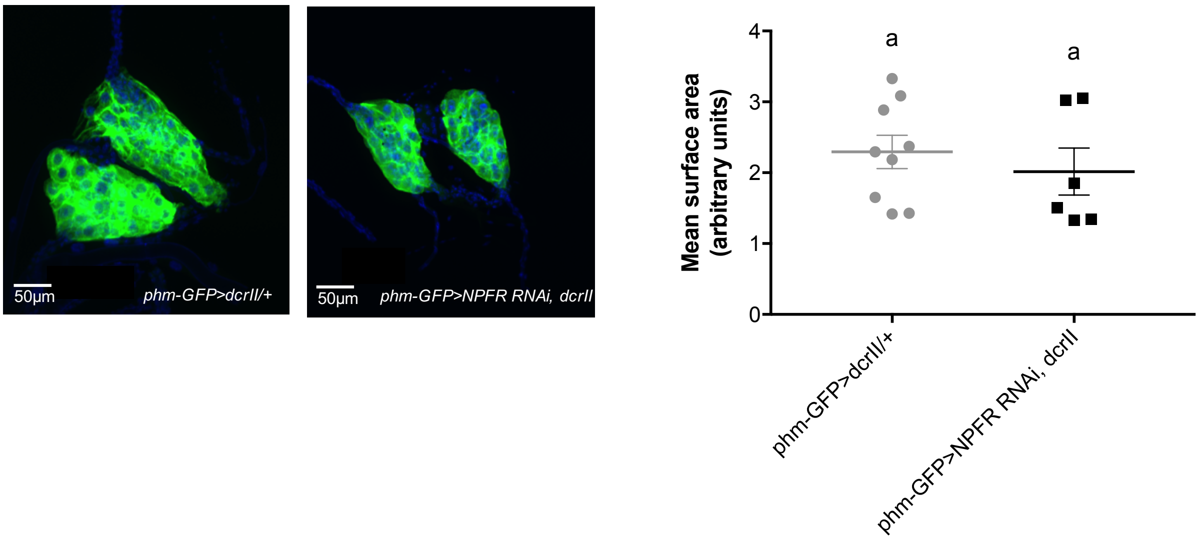


**Figure S1: Knocking down *NPFR* in the PG does not alter PG morphology**

PGs from both control and *phm-GPF>NPFR RNAi; dcrII* animals are morphologically indistinguishable. Quantification of PG size from *phm>NPFR RNAi; dcrII* and *phm>dcrII/+* animals show that there are no significant size differences. *phm-GFP; UAS-dcrII* was used to both knock down *NPFR* and to visualise PGs. Error bars represent ±1 SEM. Scale bar = 50 μm. Genotypes sharing the same letter indicate that they are statistically indistinguishable from one another ($p$ >0.05, pairwise $t$ tests). Eight-ten PGs were analysed per genotype.

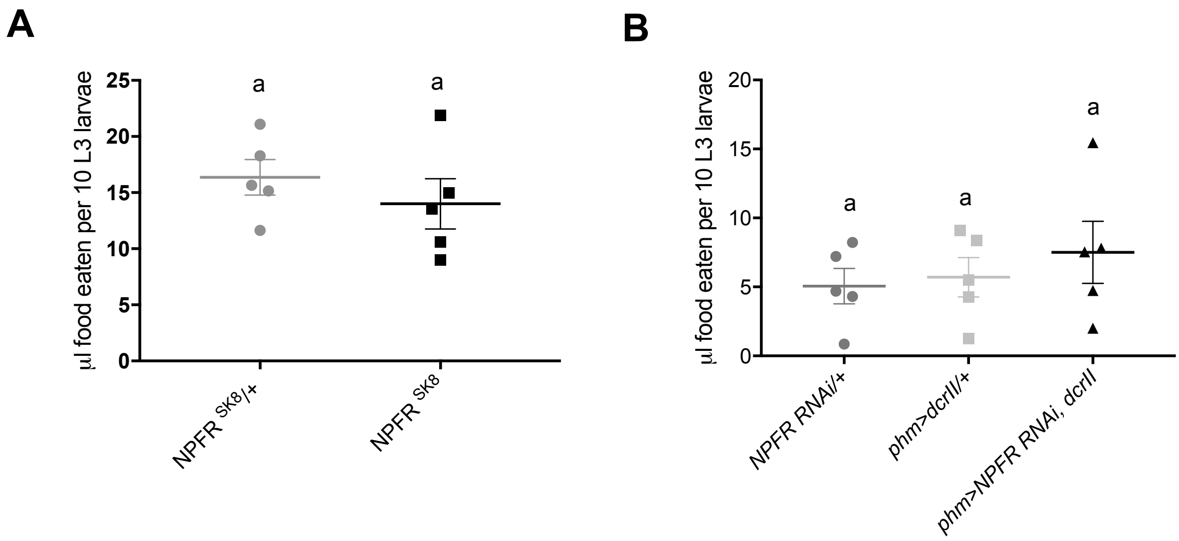


**Figure S2: Reduction of NPFR both specifically in the PG and whole-animal wide does not affect amount of food consumed**

**(A)** *NPFR*$^{SK8}$ mutants do not consume significantly more or less food compared to controls**.** Similarly, **(B)** animals where *NPFR* has been knocked down specifically in the PG do not consume significantly more or less food compared to controls. Error bars represent ±1 SEM for all graphs. Genotypes sharing the same letter indicate that they are statistically indistinguishable from one another (*p* >0.05, ANOVA and pairwise *t* tests). Each point represents a biological replicate of 10-15 newly ecdysed L3 larvae.
